# MICROCOSM FABRICATION PLATFORM FOR LIVE MICROSCOPY OF PLANT-SOIL SYSTEMS

**DOI:** 10.1101/2024.02.23.581454

**Authors:** Yangminghao Liu, Daniel Patko, Alberto Llora de le Mata, Xingshui Dong, Emma Gomez Peral, Xinhua He, Bruno Ameduri, Vincent Ladmiral, Michael P MacDonald, Lionel X Dupuy

## Abstract

Biological processes in soil pores are critical to crop nutrition and productivity, but live observations of these processes at that scale have been difficult to accomplish. To address this challenge, we have developed new techniques for the fabrication of microcosms dedicated to live imaging of the rhizosphere which incorporate the ability to control water content in transparent soil. Chambers were assembled using poly(dimethyl siloxane) (PDMS) parts fabricated by injection moulding and subsequently joined to glass slides. The control of liquid fluxes in the microcosm was achieved by syringes passing through the PDMS parts or through custom made PDMS sponges. We then tested various low refractive index materials for the fabrication of transparent soils and carried out live microscopy using Fluorescence Light Sheet microscopy. The proposed fabrication techniques are modular and enabled the construction of a wide range of experimental systems, including split chamber systems for the control of water content in soil, heterogeneous distribution of water content, monitoring of dye tracers, and live observation of plant roots. Using the techniques, we show how plant roots increase water infiltration through increased permeability of dry soil layers. This study therefore establishes that material property control and microfabrication in model rhizosphere systems can greatly enhance our understanding of plant-soil interactions.

## Introduction

Soil microscopy has been traditionally performed on thin soil sections obtained from samples embedded in resin^1^, and chemically processed to identify the location of specific organisms^2^. Using sectioning and *in situ* hybridisation techniques, it was possible to reveal the variation in microbial community composition of a Siberian tundra soil with depth^3^ or to localise bacterial species in lichens from soil crusts^4^. The technique of embedding and sectioning soil samples, though giving important insights into soil and root structure, is not suitable for live microscopy. The soil is opaque and merely allows observation of the soil surface^5^. Reconstruction of a volume dataset from sectioning techniques is time consuming because of the need to section the sample physically. Imaging techniques based on penetrating radiations are overcoming these limitations. For example, X-ray CT scanners have notably seen their resolution increase^6,7^, and phase data or high coherence beams help to resolve anatomical features from the low contrast generated by biological material^8–10^. Unfortunately, access to an X-ray CT scanner is limited and throughput very low.

Since live imaging of natural soils is limited, it has been proposed that artificial soil systems could be developed to reproduce the physical and chemical soil properties while simultaneously enabling live observations. One possibility is to use microfabrication techniques to engineer microcosms with a precisely defined arrangement of pores^11^. Various pore sizes and geometries can be imprinted on a polymer film that is in turn attached to a glass slide for direct observation. At present the behaviour of diverse microorganisms is commonly microscopically studied in such systems^12^. However, the physical obstacles imprinted are rigid, and the small pores that suits for studying microorganisms are not suitable for studying plant roots, which need to displace soil particles and create macropores that are millimetres in size^13^. Transparent soil techniques, first proposed by Downie et al.^14^, have overcome these limitations. They consist of solid particles made of transparent, low refractive index and hydrophilic materials. They can be mixed with gas and nutrient solutions to create physical conditions close to those of natural soil, leading to similar root physiology to plants grown in real soil. Live observations within such systems are possible with visible light due to a refractive index in soil particles which matches that of the soil solution^15^.

Transparent soil and microfluidic devices are ideally suited to modern light microscopy and together hold great promise for the advancement of rhizosphere science. Light microscopies are widely available instruments in biology laboratories. Modern scientific cameras and detection methods can record extremely weak signals from a range of techniques, e.g. stimulated emission depletion, fluorescence lifetime, Raman spectroscopy, or scattering^16–19^ which increase the richness of information collected from a sample. Optical sectioning has also been used to obtain data from plant roots and soil microbes *in situ* using transparent soils^20,21^.

However, to fully mimic the microcosms in which plant roots grow, environmental conditions need to be spatially controlled during experiments. Transparent soil microcosms must be compatible with the imaging stage of the microscope, a restriction which can limit the size of the samples being studied and ability to control growth conditions. Finally, there are limited soil microcosm solutions available “off the shelf” and significantly methodological development is needed prior to the running of experiments.

To address these limitations, this study presents a modular platform for the engineering of microcosm systems for application to live plant-soil microscopy. We describe the designs and protocols for the fabrication of different types of microcosm chambers and test these designs with a case study where the role of plant roots on water infiltration is demonstrated.

## Material and Method

### Injection moulding of PDMS parts of the microcosm

To achieve control fluid flux in and out of the soil (of water, nutrients and microorganisms, matching liquids, dyes etc.), microcosm chambers are assembled from glass parts through which microscopic observation and measurements are made, sandwiching poly(dimethyl siloxane) (PDMS) through which needles can be introduced for the injection and aspiration of liquid solutions and suspensions into and out from the soil (Figure 1 and Supplementary Figure 1). Moulds for the fabrication of PDMS spacers were made from three parts fabricated using a laser cutter (CO_2_ laser cutter) from methacrylate sheets (Stockline Plastics Ltd, UK). The technical drawings used for laser cutting are provided as supplementary materials at https://zenodo.org/deposit/8374546.

**Figure 1:**
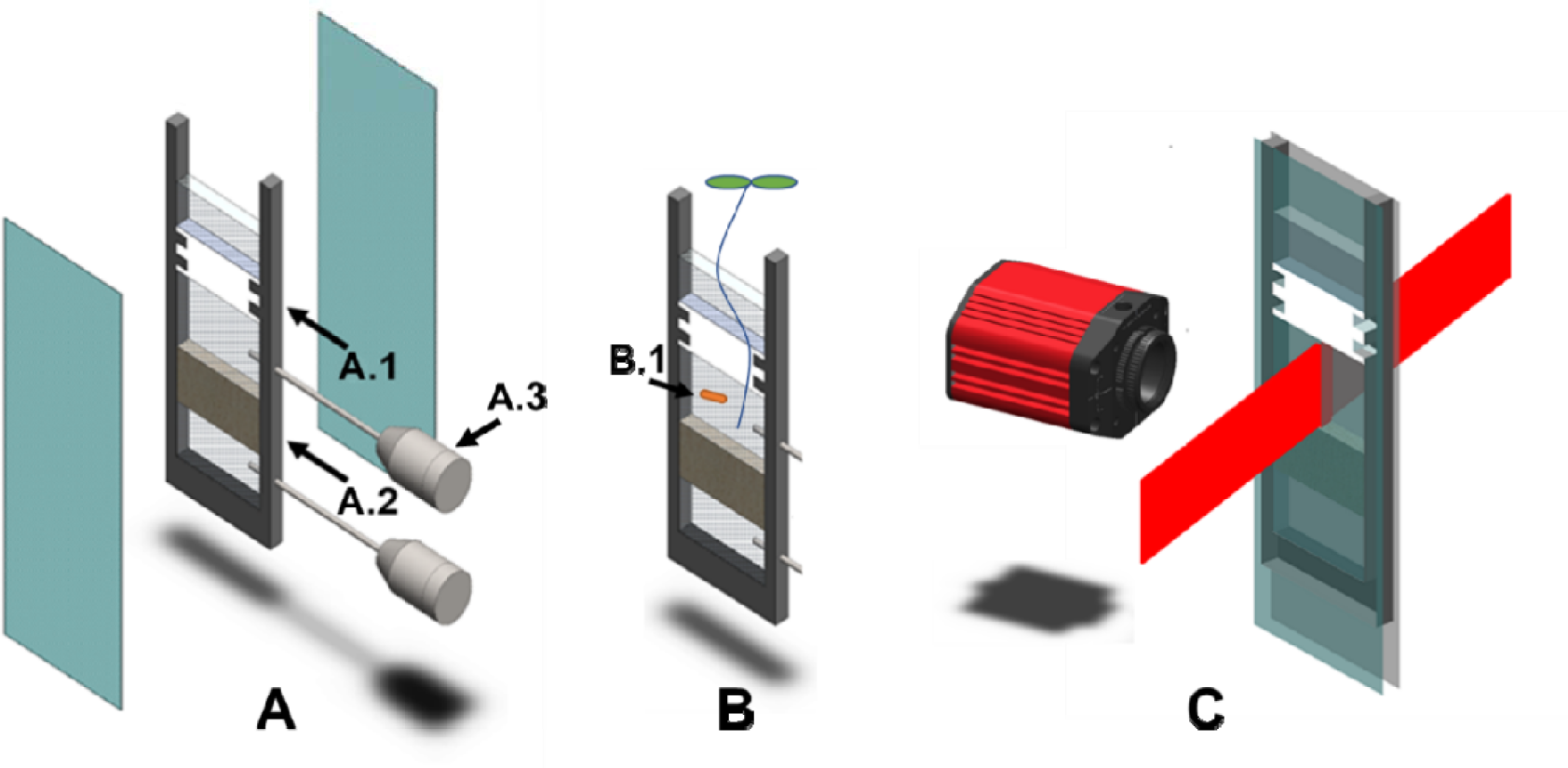
Observing environmental interactions under a microscopy requires fabrication of specialised systems that maintain growth conditions and mimics those observed in nature. (A) Microfabrication techniques can be used to engineer specialised chambers where the heterogeneity of soil properties (A.1), soil water content (A.2), and the exchange of liquid and migration of microorganisms (A.3) can be controlled. (B) Such microcosm chambers can then be filled with transparent soils, plant roots and soil microorganisms (B.1) to study biological interactions using a light sheet microscopy (C).

The central part of the mould (red parts, Figure 2 A) defines the geometry of the PDMS spacer, and the other two lateral parts (blue parts, Figure 2A) are used to seal the mould. The thickness of the methacrylate sheet defines the thickness of the transparent soil layer allowed in the microcosm. Thicknesses between 2 mm and 5 mm were tested. Three mm sheets were found to be a good compromise between the depth of imaging obtainable and the ease of filling of the transparent soil into the chamber system. The moulds were designed for the batch production of chambers. The position and size of the holes (M6) used for fastening the moulds were standardised and allowed the alignment of multiple moulds in a single operation. Bespoke geometries could be fabricated (Figure 1&2), for example to spatially constrain either the part of the chamber where the root can grow or where water can easily penetrate. The central part of the mould was either U-shaped or T-shaped. The T-shaped moulds were used to produce porous PDMS compartment for controlling water content in the microcosm. A detailed protocol for injection moulding of PDMS is provided as supplementary materials (Protocol S1).

**Figure 2:**
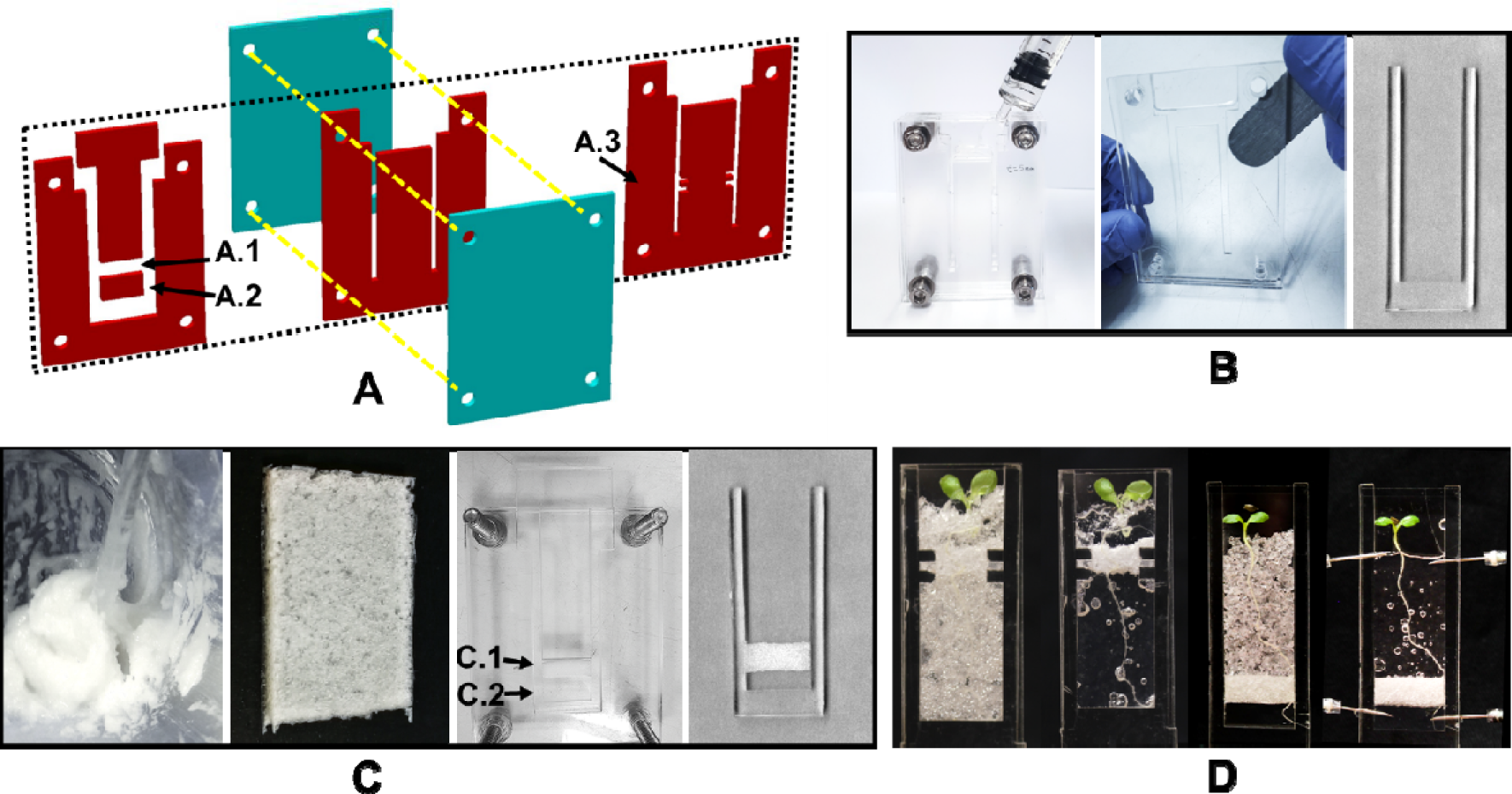
Design and fabrication of microcosm chambers. (A) Microcosms were assembled from conventional glass slides and a poly(dimethyl siloxane) (PDMS) spacer engineered by injection moulding, with the mould assembled from 3 parts with the geometry of the mould designed for controlling soil properties in the chambers. The simplest spacers had a U shape (centre) which enabled infiltration of liquids through the PMDS using syringes. Soil compartments are created using mobile parts (A.1, A.2) sliding into the central acrylic plate (left). Placing of different soil layers is aided by outgrowth (A.3). (B) PDMS injection (left) opening with a steel spatula (centre) and extraction from the central part of the mould (right) and bound to glass slides following activation of Surface using oxygen plasma. (C) Porous PDMS assemblages were built to impose a tension at the base of the soil following the principle of tention plate^46^. (D) Following the assemblage of the chamber, different soil properties can be introduced and plants are grown into the system.

### Fabrication of porous PDMS spacers for fabrication of split chambers

Porous PDMS was used to engineer compartments within the chamber. This could be used, for example, to control the water matric potential^22^. Porous PDMS was fabricated from SYLGARDTM 184 silicone elastomer base with initiator mixed at 10:1 portion and porogens made of granulated sugar (sucrose) to generate pores^23,24^. PDMS:sugar mixtures were tested in three different proportions, 2:1, 1:1 and 1:2 (v:v) respectively. Mixtures were then cured in an oven before immersion in water to dissolve the porogen. The size of pores in PDMS was controlled using milled and sieved sugar granules with three size categories: small (0.25 mm to 0.5 mm), medium (0.5 mm to 0.71 mm) and large (0.71 mm to 1.0 mm). A detailed protocol for the fabrication of porous PDMS material is supplied as supplementary information (Protocol S2).

### Assembly of chambers

Microcosm chambers were obtained by assembling the different parts described above. Glass-PDMS bonds were achieved by oxygen plasma treatment of PDMS parts, which introduced polar silanol groups (Si-OH) on the PDMS surface in turn producing Si-O-Si bonds at the expense of water molecules^25^. Plasma treatment was performed using a workbench plasma system (HPT-100, Henniker, UK) operated for 15 seconds at 100 W. Following surface treatment, glass and PDMS surfaces were joined together and pressure applied to ensure a strong bond between all adjoining surfaces. The 3D model was designed such that the PDMS spacer would fit onto standard glass microscopy slides (76 mm × 26 mm × 1 mm, VWR, UK). In order to allow to improve gas exchange and oxygen to enter the chamber, it is possible to replace either or both of the glass slides with a PDMS membrane. A detailed protocol for the assembly of glass and PDMS parts is provided as supplementary material (Protocol S3).

### Water retention of porous PDMS

To test the ability of porous PDMS to regulate water content in the transparent soil, different hydraulic properties were characterised. Porous PDMS parts were prepared with small, medium and large pore spaces using granulated sugars as porogen. Following immersion in water, porous PDMS parts were dried at room temperature before treatment by oxygen plasma for 15 s (at 100 W). Water retention curves were obtained using the tension table method (EcoTech, Bonn, Germany). Porous PDMS samples were then saturated with water before placing them in the tension plate^26^. Samples were weighed daily, and water tension was adjusted when no loss of water was recorded between two measurements. Tensions of 1 kPa, 2 kPa, 5 kPa and 10 kPa were applied and the gravimetric moisture was measured at each pressure point for each piece. All experiments were repeated three times.

### Hydraulic properties of porous PDMS

We also characterised the saturated hydraulic conductivity of porous PDMS barriers using the falling-head method. The chambers were compartmentalised with a 1 cm porous PDMS layer (Figure 2 D), with the compartment separated with small pores, medium pores and large pores in PDMS, respectively. The chambers were filled with 3.5 cm of water and a tension of constant pressure *P* was applied and the height of water in the chamber was recorded every 2 minutes. The hydraulic conductivity *K* is related to the change in the height of water in the chamber *h* through Darcy’s Law,

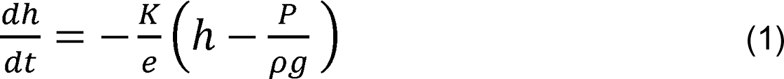

where *e* (*m*) is the thickness of the porous PDMS layer, *ρ* (*kg m*^-3^) is the density of water and *g* (*m* s^-2^) is the acceleration due to gravity. Here, the thickness of the porous PDMS layer (*e*) was measured by a ruler with an uncertainty of approximate ± 1 mm. The solution to Equation 1 is of the form

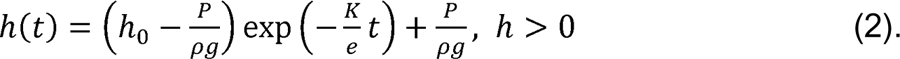

The hydraulic conductivity can therefore be estimated as (Figure 3 A)

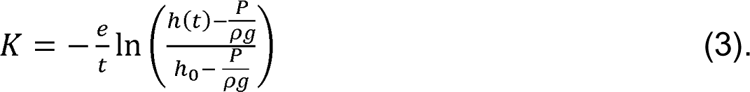

**Figure 3:**
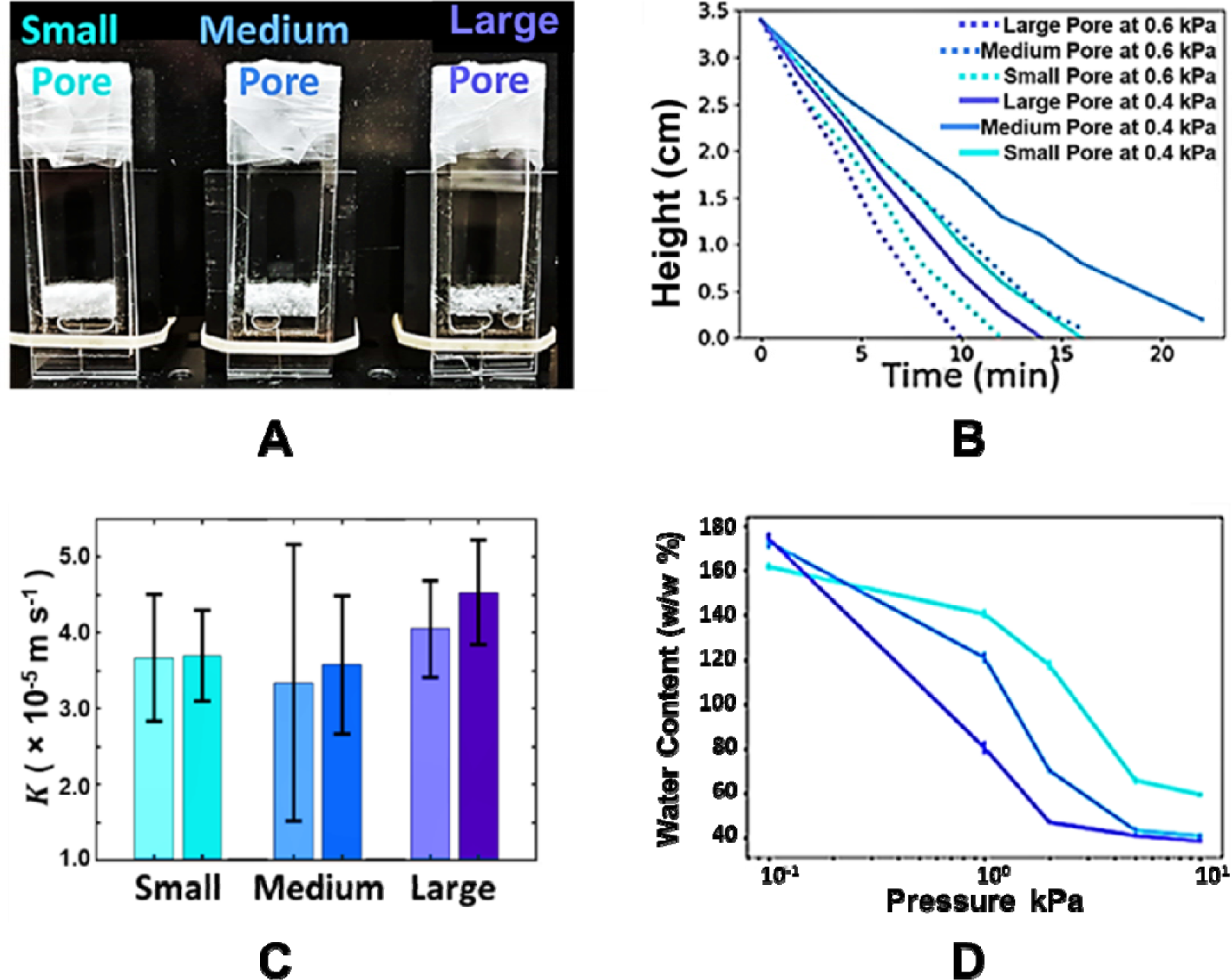
Hydraulic conductivity measurement of porous poly(dimethyl siloxane) (PDMS) in microcosm chambers. (A) Set up for falling-head measurement of the hydraulic conductivity K of the porous PDMS. A syringe is used to apply a known tension in the lower container and the height of the water column is measured at regular time interval. (B) Results show that the flux of water through porous PDMS occurs faster with small pores than with medium pores, but slower than with large pores. The hydraulic conductivity is quantitatively shown for different pore sizes (C). The small pores have the greatest water retention (D) which makes it suitable for regulating water content in transparent soil.

All experiments were repeated 6 times.

### Low refractive index materials and their suitability for growing and live imaging of roots

Before plants can be grown in microcosm chambers, water and various types of transparent soil substrates were introduced. The materials tested for the fabrication of transparent soils consisted of Nafion™ particles (IonPower, USA), fluorinated ethylene propylene (FEP) particles (Holscot, UK), hydrogel beads ^27^, and Forblue™ particles (tube particles of two types of :outside diameter: 3 and 0.5 mm, SELEMION^TM^ tubes, AGC, Japan). Nafion™ particles (0.25 to 1.25 mm) were prepared according to previous work^14^. FEP pellets (approximately 2 mm) were sieved from raw materials. To assess the effect of substrates on growth, agricultural soil (sandy loam, 71% sand, 19% silt and 10% clay, Bullion field at the James Hutton Institute, Dundee, 56°27’34” N 3°4’21”W) and agar medium (wt 1.5%, Sigma A1296-100G) were also used. A bio-compatibility assay was implemented to test the suitability of these various materials. Seeds of lettuce (*Lactuca sativa* “all year round”, Sutton Seeds, UK) were surface sterilised by washing in 10% bleach for 15 minutes followed by thorough rinsing with sterile dH_2_O. Sterilised seeds were placed in agar on a petri dish and incubated at 21 °C in a growth facility for germination. 7 ml of the different processed substrates were poured in glass tubes (16 mm × 100 mm, OD × length) and saturated in water, and seedlings with similar size were grown in 6 replicate experiments.

A second experiment was performed to assess the image quality obtained when using the above-mentioned substrates. The microcosm chambers were first autoclave sterilised. Seedlings were germinated as described above and transferred under sterile laminar air flow. Seedlings were grown for 2 days at 21°C with 16 hours light and 8 hours dark period (Figure 2.5). A colloidal suspension (Percoll^®^, GE Healthcare, UK) at 80% v/v in dH_2_O was used for refractive index matching of sulforrhodamine-B (Sigma-Aldrich, UK) stained at 1 µg/100 ml concentration. When imaging through Nafion™ and Forblue™ particles, light scattering was acquired using 633 nm illumination and recorded without a filter. The stained Nafion™ and Forblue™ particles were illuminated at 561 nm and recorded with a long pass filter (cut-on wavelength: 570 nm, Thorlabs).

### Light sheet microscopy

Images were acquired using a custom-made Light Sheet Fluorescence Microscopy (LSFM) developed by Liu et al.^21^. A four wavelength-laser source (488 nm, 514 nm, 561 nm and 633 nm, Vortran Versalase, Laser 2000 Ltd, UK) was used to create a thin sheet of light to illuminate the sample. Two counterpropagating light sheets sectioned the sample, one from each of two sides of the chamber. A microscopy objective (5X, Mitutoyo, Japan) and a digital camera (Orca flash 4, Hamamatsu, Japan) were perpendicularly placed to the light sheet to capture light emitted from the sample, either through scattering or fluorescence, at a maximum dynamic range (16 bits of greyscale). To isolate the fluorescence signal, the imaging arm was respectively fitted with long pass filters of 530 nm, 570 nm, 645 nm and 665 nm (FGL530, FGL570, FGL645, FGL665, Thorlabs, UK).

### Case Study: Effect of root growth on water infiltration

Chambers were created using 4 mm thick PDMS spacers. Wet soil was added to the top and bottom. An 8 mm barrier of dry soil was created using the same Nafion™ particles that were oven dried at 100°C overnight. Drying of Nafion™ causes the water contact angle to change from 25° (hydrophilic) to 105° (hydrophobic)^28^. Dry soil is difficult to be rewetted without an acidification process ^14^, and hence, placing the dry soil between two wet soil layers forms a hydrophobic layer. A winter wheat seedling (*Triticum aestivum* var. Filon) and a tracer dye were placed in the upper wet layer. In the control samples only the tracer dye was added, but preparation was otherwise the same. The tracer dye is composed of a concentration of 0.1 mL of red dye (Red food colorant, Vahiné, Spain) in a volume of 1 mL of sterile water. To accurately evaluate the concentration of the dye, an imaging analysis method was developed. Chambers filled with tracer dye solutions concentration 0, 0.025, 0.05, 1, 2, 4, 6, 8, 10, 12, 14, 16, 18 or 20 × 10^-4^ (volume/volume) were imaged by a colour camera (EOS 100, Cannon) in front of a backlight consisting of a white monitor screen. The images acquired by the camera (Figure 5A and C), cropped and transformed to HSV coordinate system (Hue, Saturation, Brightness or Value) (Figure 5A right). A piecewise function relating the Saturation (S) and Brightness (V) to the dye concentration was then established,

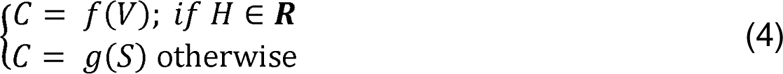

Where C is the concentration of the dye solution and ***R*** is the hue range of the calibrated red colour. The algorithm was implemented in MATLAB (MathWorks Inc, USA) and subsequently used to estimate the concentration of the dye in the chamber. To quantify the infiltration of the tracer dye through the microcosm, a quantitative measurement of the saturation of images taken daily with a camera (EOS 100, Cannon) was performed. An RGB (Red-Green-Blue) image was obtained and the permeability (k) was calculated using the following formula:

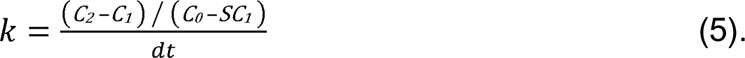

The concentration in the upper (*C_0_*) and lower (*C_1_*) soil layers were measured during the first day and the lower region (*C_2_*) was imaged during the following days. dt is the time interval between the measurements *of C_1_* and *C_2_*.

## Results and discussion

### Microfabrication techniques allow to engineer growth microcosms for visualizing plant and soil microorganisms

Using the fabrication techniques outlined above, we have constructed and tested various chamber designs (Figure 2D). The microcosm chambers provided access for the preparation of custom microcosms for confocal imaging, light-sheet imaging and for optical manipulation of plant roots. This was aided by the maintenance of a constant distance between the two glass parts as set by the U-shapes produced by PDMS spacers. The PDMS spacer allowed easy perforation by needles while maintaining a watertight seal during experiments (Figure S1). The covalent bonds (Si-O-Si) between PDMS and glass were durable and maintained after several autoclave treatments such that the chambers were reusable after cleaning. Importantly, the design allowed for the control of fluids inside the microcosms created (Supplementary Video S1), including through the porous PDMS compartment.

Similar materials and fabrication techniques have been used in other studies. For instance, PDMS has typically been used to engineer structures at a smaller scale^29–31^ or cantilever systems (measurement of forces), when forming a nanoporous interface (exudate sampling), and to introduce pores. These systems are less easily transferred to other labs because of the need for specialised equipment and facilities as well as the knowhow to operate them. Simpler fabrication techniques such as that proposed by EcoFAB have been demonstrated^32,33^. With our framework we showed that such fabrication techniques can be expanded to combine features that allow for the control of water in the microcosms created (fluxes, content and heterogeneity), which is critical in the study of plant responses to their environment. Fabrication of such chambers could be further facilitated using 3D printing, which can now rapidly build PDMS structures of centimetre to millimetre size^21,32–34^.

### Engineered porous barriers enable control of water flow and soil water content in the chamber

Various porogen compositions were tested to create tuneable porous barriers in the system (Figure 2D). PDMS with low porogen content (1:2, v:v) did not show water holding abilities nor any form of permeability to air. This was due to lack of connectivity between sugar particles, which simultaneously prevented the removal of sugar from the inner parts of the PDMS whilst not creating the interconnected pore spaces needed for fluid transport. Improvements were achieved with PDMS mixtures with increased sugar content (1:1, v:v), but a complete removal of sugar was not possible either. PDMS with high sugar content (2:1, v:v) enabled a complete removal of sugar and permeability of the porous material through interconnected pore spaces. The hydraulic conductivity of the porous PDMS compartment was tested using the falling-head method (Figure 3 B&C). The results indicated the hydraulic conductivity of PDMS under a tension of 0.4 kPa or 0.6 kPa was 3.7 ± 0.8 × 10^-5^ ms^-1^ or 3.7 ± 0.6 × 10^-5^ ms^-1^ with small pores (>0.25 but <0.50 mm), 3.3 ± 1.8 × 10^-5^ ms^-1^ or 3.6 ± 0.9 × 10^-5^ ms^-1^ with medium pores (>0.50 but <0.71 mm), and 4.1 ± 0.6 × 10^-6^ m s^-1^ or 4.5 ± 0.7 × 10^-5^ m s^-1^ with large pores (>0.71 mm but <1.00 mm), respectively. The flux of water through PDMS occurs faster with small pores than with medium pores but slower than with large pores.

The water retention of porous PDMS with the small pores was greatest compared with the medium pores and large pores, particularly when the tension ranged from 0 kPa to 5 kPa. When the tension reached 10 kPa, the water in the porous PDMS reduced significantly (Figure 3D). Application of a tension of 0.4 kPa removed all water from the top part of the chamber. The curves showing the flux of water through porous PDMS showed that pore size consistently affected the transport of water.

Overall, the results of the hydraulic characterisation of porous PDMS layers were as expected. Smaller pore sizes increased water retention and reduced the permeability of the barrier. The hydraulic conductivity of PDMS was moderate and similar to that of sandy loam soils^35^. The water retention, enabled by the porous PDMS, was similar to sand^36^. This system may not be suitable to control water content in a natural soil (only wet conditions), but is adequate when using artificial soils such as those made of FEP^37^ or Nafion™^14^, which have similar water retention properties.

### Suitability of materials for the fabrication of transparent soils

Light sheet image data were acquired from a range of substrate types (Fig 4A). Nafion™-based transparent soil stained with sulforhodamine-B produced images where both root and particles could be visualised with great clarity (Figure 4A). Images of similar quality were obtained with Forblue™ based transparent soil. Images acquired from roots grown in FEP based transparent soil showed the lowest quality of those investigated here. FEP is less transparent than other materials tested and image noise from the Nafion™ soil is most likely due to scattering from within the FEP itself, whilst also coming from Fresnel reflections at air gaps between the FEP and water due to the hydrophobic behaviour of surfaces. Soils made of hydrogel beads were transparent and image quality obtained with the system showed similar properties to those obtained in Nafion™- and Forblue™-based transparent soil.

**Figure 4.**
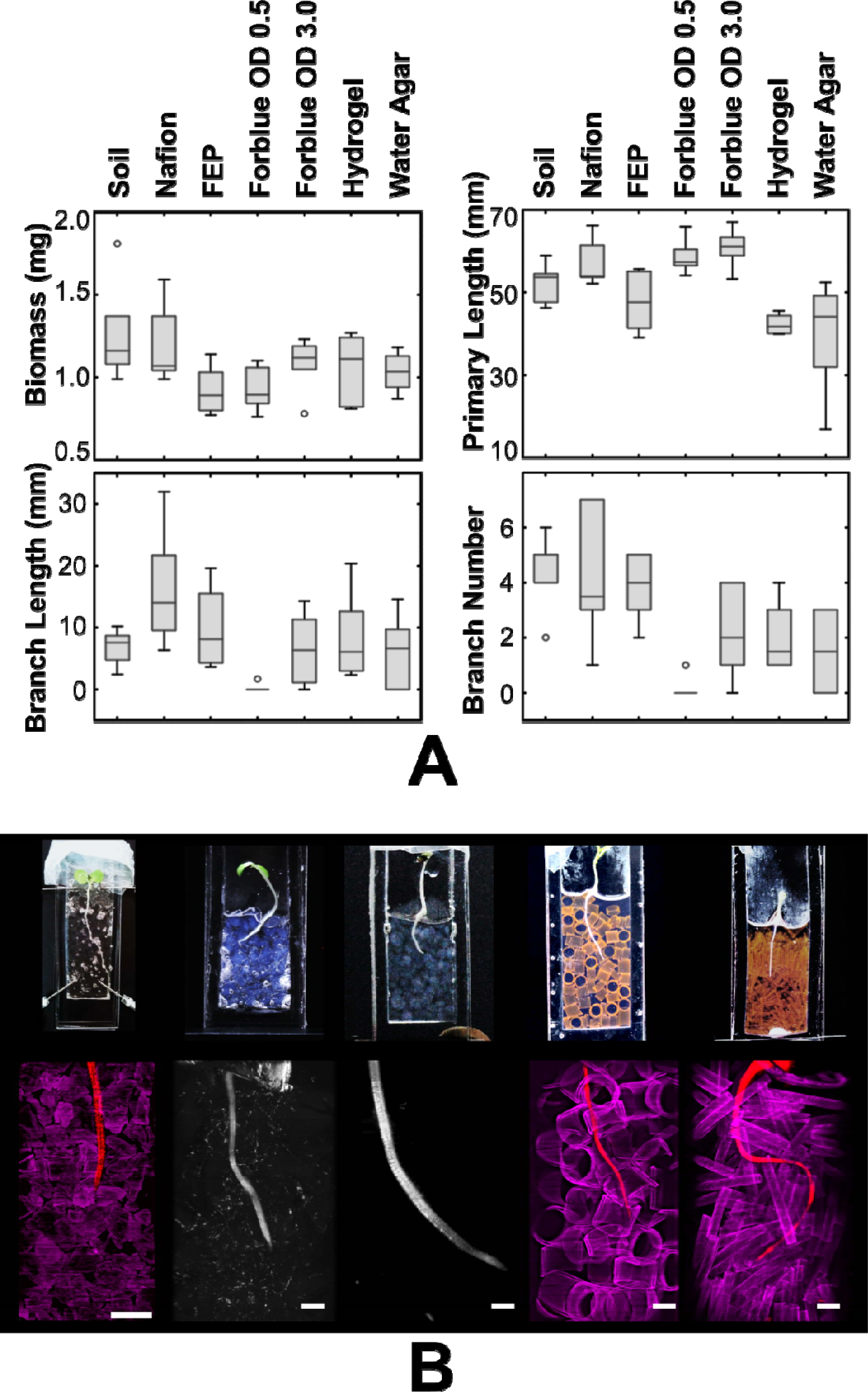
Testing of different materials for growth and live imaging through artificial soils. (A) Growth of lettuce in the five transparent soils compared to growth in a natural soil and in two agar media (hydrogel beads and plain water agar). (B) Live imaging of lettuce seedling grown in transparent soils made of five low refractive index materials. Top left to right: lettuce seedlings in Nafion™ particles, fluorinated ethylene propylene (FEP) particles, hydrogel particles, sulforrhodamine-B stained Forblue™ particles (3 mm outside diameter), and sulforrhodamine-B stained Forblue™ particles (0.5 mm outside diameter). Bottom left to right: maximum projection images of lettuce roots (red) in sulforrhodamine-B stained Nafion™ (magenta), Llettuce root (gray) in FEP particles (grey), lettuce root (grey) in hydrogel particles (grey but barely imaged), lettuce root (red) in sulforrhodamine-B stained Forblue™ particles of 3 mm OD (magenta), lettuce root (red) in sulforrhodamine-B stained Forblue™ particles of 0.5 mm OD (magenta). All scale bars represent 2 mm.

**Figure 5.**
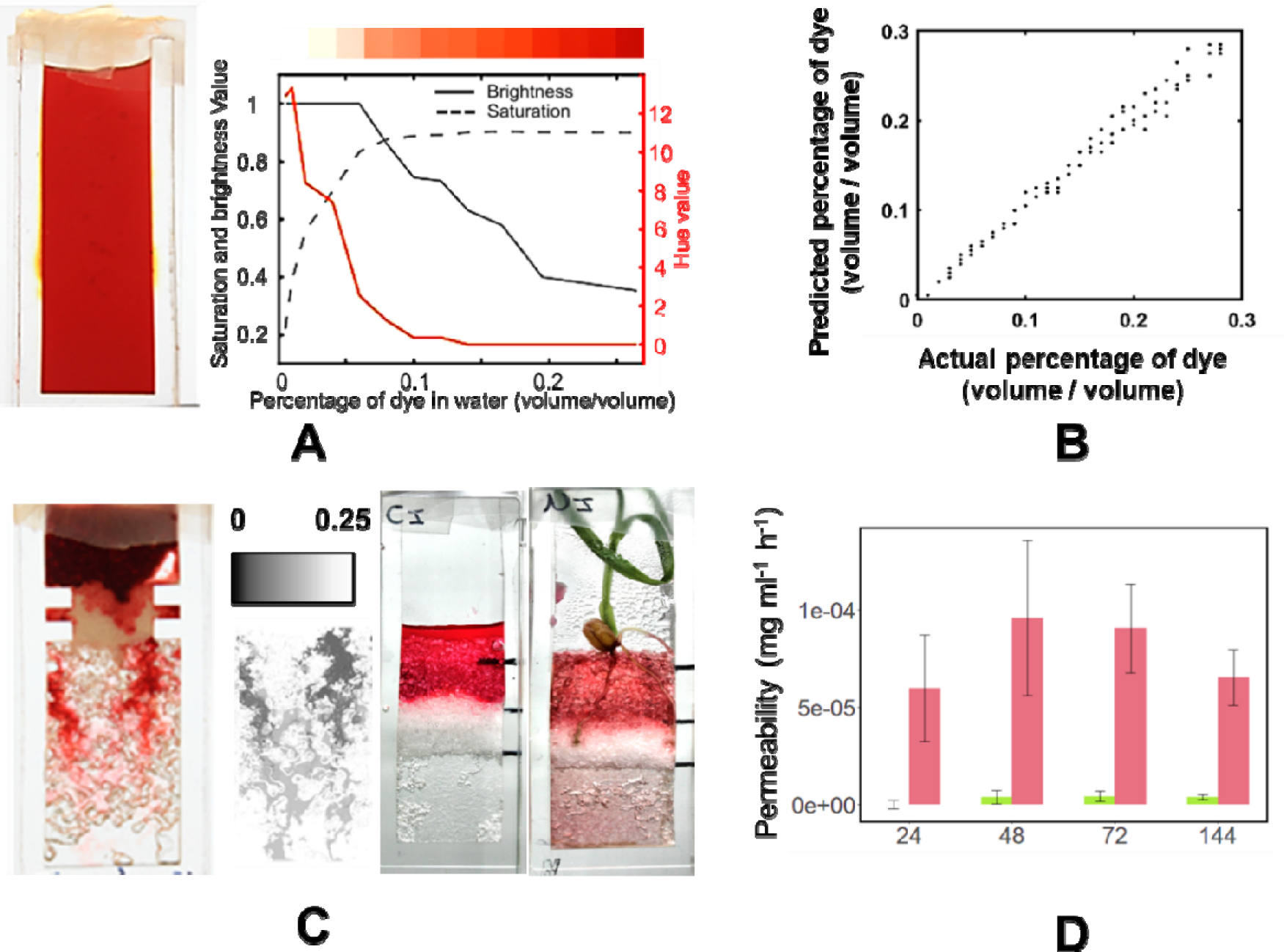
Application of microcosm chambers to the characterisation of water transport through dry soil layers. (A, left) Photo of the dye-water solution at 0.18 % (volume/volume). (A, right) Relationship of dye concentration and recorded HSV values from the photo. (B) Correlations between experimental and predicted dye concentration. (C) The dye solution infiltrated through the dry soil layer (left) can then be determined through the permeability of the barrier (equation 4). Results show the root facilitated the transport of water through the dry barrier (D).

Overall, plants had a higher dry weight in soil and in Nafion™ than with other artificial mediums (Figure 4B). The primary root length was longer in Nafion™ particles and Forblue™ particles. However, more secondary roots were seen in agricultural soil. Roots grown in Forblue™ particles with 0.5 mm diameter were frequently damaged during extraction. Plants grown in soil, Nafion™, large Forblue™ particles, and hydrogel beads had slightly higher dry weight than those grown in agar. More secondary roots were observed in plants grown in soil, Nafion™, FEP and large Forblue™ particles.

In general, results showed that plant growth was better modelled when growth was in artificial soils than in a gel medium, which confirmed previous experiments^14,27^. No plants showed signs of stress and overall, all materials would be considered as biocompatible. Most materials showed capability to provide high quality live images of the rhizosphere. Use of low cost, low refractive index polymers such as FEP produced lower quality data, but the recent methods for functionalising FEP ^37^ indicated potential for significant improvement of the technique.

The results in this study confirm it is possible to construct centimetre-scale microcosms for plants with the ability to measure root development up to one week. The physical model constructed from transparent and soil-like substrates is able to mimic the heterogeneous nature of soil, something which is the key to forming boundaries in physiochemical gradients, root exudates, and microbial communities^38^. Indeed, spatial and temporal associations in their rhizosphere were observed in our microcosms chambers, where root systems developed more lateral roots^14^, bacteria cells formed collective movements^20,21^, and rhizosphere acidification was from root exudate^37^. Spatial and temporal dynamics in transparent soil present a smaller but more dynamic rhizosphere when compared to a gel-based system, where homogenous diffusion might lead to overestimated rhizosphere ranges^39^ and other non-physiologically representative behaviours.

### Image analysis demonstrates how root growth affects the permeability of dry soil layers

It was possible to create a heterogeneous water distribution by adding a hydrophobic barrier to an imaging chamber (dry Nafion™ particle layer as thin as 3 mm in height). The glass window of the microcosm chamber allowed measurements based on quantitative imaging methods developed previously. Here, measurements were made in the HSV colour space and allowed estimation of the permeability of dry soil layers to a dye tracer (Figure 5A & B). The methods showed there was a nearly linear relationship between dye concentration and either the brightness (in the range [0.00, 0.06]) and saturation (in the range [0.06, 0.20]). We applied the method to quantify the infiltration of water through dry soil with and without plants (Figure 5C). Results showed that the growth of plant roots significantly increased the permeability of the dry soil barrier even though water uptake resulted in a reduced pressure head at the top of the dry soil layer (Figure 5D). A Kruskal-Wallis rank sum test was used to confirm that the effect of the root was statistically significant (χ^2^ = 24.237, *df* = 1, *P*-value= 8.5 × 10^-7^). After 24 hours introduction of the tracer dye, coloration was observed on the sample containing roots with a mean permeability of 6.0 x 10^-5^ mg ml^-1^ h^-1^ in the dry soil. In the following 72 hours, the permeability increased slightly, probably due to the slow rewetting of the dry soil but remained to similar order of magnitude. In contrast, the coloration of the bottom soil layer in control treatments was negligible and permeability was not significant.

These results showed that microcosm experiments had significantly potential for studying water fluxes in the rhizosphere. Hydrologists have long established that plants modify water movements in soil through preferential flow^40,41^, which limits runoff and its associated soil erosion. However, different types of plants can have varying effects on water infiltration^42,43^, and the role of biological activity (roots and microorganisms secretion, microbial growth) in these processes is yet to be clarified^44,45^.

## Conclusion

We have conducted experiments to refine and streamline the preparation of soil microcosm systems designed for live, *in situ* imaging of root soil interactions. Our approach involves the rapid prototyping and reproduction of chamber designs in a laboratory setting, employing techniques such as 3D printing, laser cutting, and cast molding of parts. Intensive testing of the system has revealed its remarkable tunability, especially with regard to environmental conditions and the control of water content, enhancing the reproducibility and reliability of experiments. Furthermore, our exploration of various materials has identified several as suitable for fabricating transparent soil substrates. In specific experiments, we have introduced a heterogeneous distribution of soil water content and employed tracer dyes to quantify water infiltration. This study demonstrates that our system can be applied to effectively address current knowledge gaps in understanding plant-soil interactions.

## Acknowledgement

This work was funded by the European Research Council (ERC) under the European Union’s Horizon 2020 research and innovation programme (Grant agreement No. 647857-SENSOILS). We also acknowledge the funding from the Spanish Ministry of Science and Innovation (MICINN) under de project MICROCROWD (PID2020-112950RR-I00). We thank Dr Masahiro Oh from AGC Inc for the supply of Forblue™ free samples.

## Supplementary Materials

**Figure S1.**
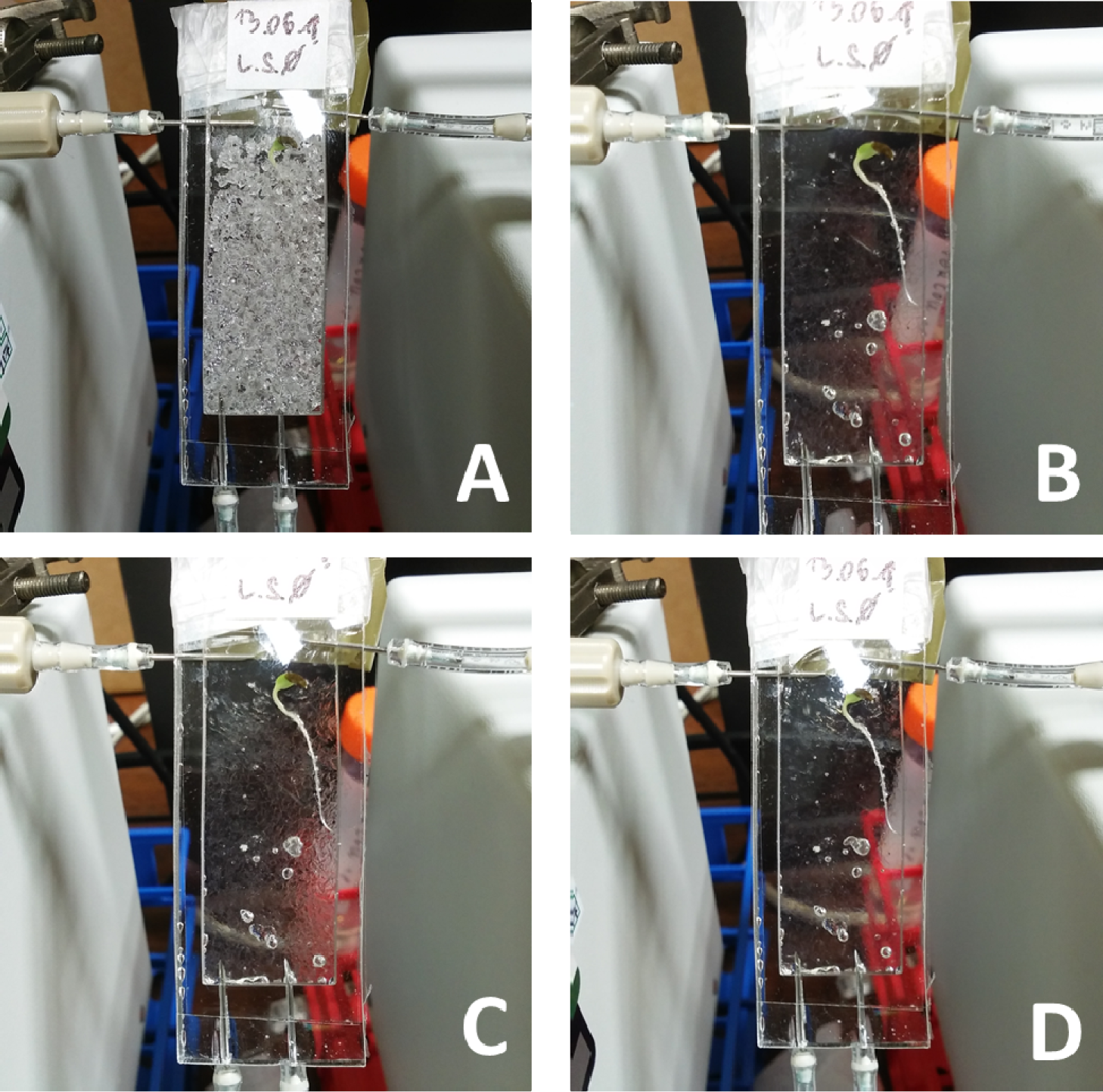
Testing of the flow of liquid through the transparent soil chambers. A lettuce seedling grown into the chamber (A). Two liquids are successfully perfused into the soil. (B) The matching liquid (80% Percoll ® dispertion) is firstly introduced, and visibility of the background through the sample indicates good circulation of the liquid. Water is then introduced 10 minutes later (C). The experiment was reproduced multiple times in using the same samples, indicating the repeatability of the system. After 10 iterations, refractive index matching is still successfully obtained (D).

## Technical drawings files

*Technical drawing S1*

PDMS cast mould Bottom and Cover

*Technical drawing S2*

PDMS cast mould version A Part1

*Technical drawing S3*

PDMS cast mould version A Part2

*Technical drawing S4*

PDMS cast mould version B

All files are available in dxf format at https://zenodo.org/deposit/8374546

## Protocols

### Protocol S1

Fabrication of PDMS used for the assembly of microcosms.

1. SYLGARDTM 184 Silicone Elastomer Base and the curing agent hardener mixed at 10:1 portion.
2. Assemble the U-shape moulds with fasteners (M6 bolts, washers and nuts).
3. Injection of the PDMS liquid into the U-shapes mould using a syringe. A PDMS joint with 3 mm width will use a volume of approximately 1.8 ml (Figure 2 B).
4. Apply vacuum the mould in upright position for 60 min (at approximately 70 kPa), followed by 15 min at atmospheric pressure. Reiterate cycle until total removal of air bubbles into the mould.
5. Wrap the mould in aluminium foil and cure in upright position in an oven at 70°C. The time for curing U-shape mould of 3 mm width is around 1h 30.
6. Cool down the sample for a few minutes before extraction of PDMS spacer from mould. Remove fasteners and open the mould using a thin steel lever (Figure 2 B).
7. Use a scalpel to smooth the edge of the extracted PDMS spacer.

### Protocol S2

The fabrication process of PDMS spacer with porous PDMS compartment is outlined below:

1. Mix and stir the PDMS and sugar until homogenous (the white mixture in Figure 2 C).
2. Fill the mixture in the mould assembled a Figure 2 A, without the centre T-shape.
3. Assemble the mould and cure the porous PDMS in an oven at 70°C for 2 hours.
4. Open the mould, extracted the cured porous PDMS piece, and immersed in water for 2 days to dissolve particles. Rinse the porous PDMS piece and dry in room temperature (Figure 2C).
5. The U shape PDMS spacer is place back into a T shape moulds (Figure 2A and C) the porous PDMS mixture is subsequently added at the correct height (Figure 2 A.1 and C.1).
6. The porous PDMS part is then cured in oven at 70°C for 2 hours before extraction while in contact with the U shape PDMS to allow bounding of both parts.
7. the PDMS assembly is then immersed in water for two days and air dried at room temperature.

### Protocol S3

The detailed protocol is as follows:

1. The PDMS spacer is placed in 3D printed assembly.
2. The glass slide and the assembly tool with PDMS spacer was placed in oxygen plasma system for 15 seconds at 100 W.
3. The glass and PDMS surface exposed to plasma were aligned and joined using the assembly tool. Gentle pressure is applied to the glass to obtain homogeneous contact with PDMS.
4. The semi assembled chamber is turned upside down and placed in the plasma system, together with a second glass slide. Plasma treatment is subsequently applied for 15 seconds at 100W.
5. The glass and PDMS surface exposed to plasma are aligned and joined using the assembly tool. Gentle pressure is applied to the glass to obtain homogeneous contact with PDMS.
6. The chamber is extracted from the assembly tool.

## Reference list

1 R. Tippkötter and K. Ritz, Geoderma, 1996, 69, 31–57.

2 N. Nunan, K. Ritz, D. Crabb, K. Harris, K. Wu, J. W. Crawford and I. M. Young, FEMS Microbiology Ecology, 2001, 37, 67–77.

3 S. Kobabe, D. Wagner and E.-M. Pfeiffer, FEMS Microbiology Ecology, 2004, 50, 13–23.

4 L. Muggia, B. Klug, G. Berg and M. Grube, Applied Soil Ecology, 2013, 68, 20–25.

5 K. Ritz and I. M. Young, Mycologist, 2004, 18, 52–59.

6 A. van Veelen, N. Koebernick, C. S. Scotson, D. McKay-Fletcher, T. Huthwelker, C. N. Borca, J. F. W. Mosselmans and T. Roose, New Phytologist, 2020, 225, 1476–1490.

7 H. Zhou, W. R. Whalley, M. J. Hawkesford, R. W. Ashton, B. Atkinson, J. A. Atkinson, C. J. Sturrock, M. J. Bennett and S. J. Mooney, Journal of Experimental Botany, 2021, 72, 747–756.

8 N. Kunishima, Y. Takeda, R. Hirose, D. Kalasová, J. Šalplachta and K. Omote, Plant Methods, 2020, 16, 7.

9 M. Sedighi Gilani, M. N. Boone, K. Mader and F. W. M. R. Schwarze, Journal of Structural Biology, 2014, 187, 149–157.

10 P. Verboven, E. Herremans, L. Helfen, Q. T. Ho, M. Abera, T. Baumbach, M. Wevers and B. M. Nicolaï, The Plant Journal, 2015, 81, 169–182.

11 C. E. Stanley, G. Grossmann, X. C. i Solvas and A. J. deMello, Lab Chip, 2016, 16, 228–241.

12 P. M. Mafla-Endara, C. Arellano-Caicedo, K. Aleklett, M. Pucetaite, P. Ohlsson and E. C. Hammer, Commun Biol, 2021, 4, 1–12.

13 J. Aufrecht, M. Khalid, C. L. Walton, K. Tate, J. F. Cahill and S. T. Retterer, Lab Chip, 2022, 22, 954–963.

14 H. Downie, N. Holden, W. Otten, A. J. Spiers, T. A. Valentine and L. X. Dupuy, PLOS ONE, 2012, 7, e44276.

15 F. E. O’Callaghan, R. A. Braga, R. Neilson, S. A. MacFarlane and L. X. Dupuy, Sci Rep, 2018, 8, 1440.

16 W. Becker, Journal of Microscopy, 2012, 247, 119–136.

17 C. W. Freudiger, W. Min, B. G. Saar, S. Lu, G. R. Holtom, C. He, J. C. Tsai, J. X. Kang and X. S. Xie, Science, 2008, 322, 1857–1861.

18 S. W. Hell and J. Wichmann, Opt. Lett., OL, 1994, 19, 780–782.

19 Z. Yang, H. Downie, E. Rozbicki, L. X. Dupuy and M. P. MacDonald, Opt. Express, OE, 2013, 21, 16239–16247.

20 I. C. Engelhardt, D. Patko, Y. Liu, M. Mimault, G. de las Heras Martinez, T. S. George, M. MacDonald, M. Ptashnyk, T. Sukhodub, N. R. Stanley-Wall, N. Holden, T. J. Daniell and L. X. Dupuy, ISME J, 2022, 16, 2337–2347.

21 Y. Liu, D. Patko, I. Engelhardt, T. S. George, N. R. Stanley-Wall, V. Ladmiral, B. Ameduri, T. J. Daniell, N. Holden, M. P. MacDonald and L. X. Dupuy, Proceedings of the National Academy of Sciences, 2021, 118, e2109176118.

22 W. R. Whalley, E. S. Ober and M. Jenkins, Journal of Experimental Botany, 2013, 64, 3951–3963.

23 K. J. Cha and D. S. Kim, Biomed Microdevices, 2011, 13, 877–883.

24 S.-J. Choi, T.-H. Kwon, H. Im, D.-I. Moon, D. J. Baek, M.-L. Seol, J. P. Duarte and Y.-K. Choi, ACS Appl. Mater. Interfaces, 2011, 3, 4552–4556.

25 S. Bhattacharya, A. Datta, J. M. Berg and S. Gangopadhyay, Journal of Microelectromechanical Systems, 2005, 14, 590–597.

26 R. R. Bruce and A. Klute, Soil Science Society of America Journal, 1963, 27, 18–21.

27 L. Ma, Y. Shi, O. Siemianowski, B. Yuan, T. K. Egner, S. V. Mirnezami, K. R. Lind, B. Ganapathysubramanian, V. Venditti and L. Cademartiri, Proceedings of the National Academy of Sciences, 2019, 116, 11063–11068.

28 Wetting and Absorption of Water Drops on Nafion™ Films | Langmuir, https://pubs.acs.org/doi/10.1021/la800799a, (accessed January 25, 2024).

29 K. Ozoe, H. Hida, I. Kanno, T. Higashiyama and M. Notaguchi, in 2015 28th IEEE International Conference on Micro Electro Mechanical Systems (MEMS), 2015, pp. 702–705.

30 D. E. W. Patabadige, L. J. Millet, J. A. Aufrecht, P. G. Shankles, R. F. Standaert, S. T. Retterer and M. J. Doktycz, Sci Rep, 2019, 9, 10272.

31 S. R. Lockery, K. J. Lawton, J. C. Doll, S. Faumont, S. M. Coulthard, T. R. Thiele, N. Chronis, K. E. McCormick, M. B. Goodman and B. L. Pruitt, Journal of Neurophysiology, 2008, 99, 3136–3143.

32 J. Gao, J. Sasse, K. M. Lewald, K. Zhalnina, L. T. Cornmesser, T. A. Duncombe, Y. Yoshikuni, J. P. Vogel, M. K. Firestone and T. R. Northen, JoVE (Journal of Visualized Experiments), 2018, e57170.

33 J. Sasse, J. Kant, B. J. Cole, A. P. Klein, B. Arsova, P. Schlaepfer, J. Gao, K. Lewald, K. Zhalnina, S. Kosina, B. P. Bowen, D. Treen, J. Vogel, A. Visel, M. Watt, J. L. Dangl and T. R. Northen, New Phytologist, 2019, 222, 1149–1160.

34 S. Ge, L. X. Dupuy and M. P. MacDonald, Plant Soil, 2021, 468, 475–489.

35 D. M. Allen, A. J. Cannon, M. W. Toews and J. Scibek, Water Resources Research, 2010, 46, W00F03.

36 W. Durner, in Encyclopedia of Soils in the Environment (Second Edition), eds. M. J. Goss and M. Oliver, Academic Press, Oxford, 2023, pp. 168–186.

37 D. Patko, Q. Yang, Y. Liu, P. Falireas, B. Briou, B. V. Tawade, T. S. George, T. J. Daniell, M. P. MacDonald, V. Ladmiral, B. Ameduri and L. X. Dupuy, Plant Soil, , DOI:10.1007/s11104-023-06151-y.

38 Y. Kuzyakov and B. S. Razavi, Soil Biology and Biochemistry, 2019, 135, 343–360.

39 M. O. Yee, P. Kim, Y. Li, A. K. Singh, T. R. Northen and R. Chakraborty, Frontiers in Microbiology.

40 J. Bouma, Agricultural Water Management, 1981, 3, 235–250.

41 M. Flury, H. Flühler, W. A. Jury and J. Leuenberger, Water Resources Research, 1994, 30, 1945–1954.

42 V. H. Durán Zuazo and C. R. Rodríguez Pleguezuelo, Agron. Sustain. Dev., 2008, 28, 65–86.

43 L. Jačka, A. Walmsley, M. Kovář and J. Frouz, Geoderma, 2021, 403, 115372.

44 M. Naveed, M. A. Ahmed, P. Benard, L. K. Brown, T. S. George, A. G. Bengough, T. Roose, N. Koebernick and P. D. Hallett, Plant Soil, 2019, 437, 65– 81.

45 D. Or, S. Phutane and A. Dechesne, Vadose Zone Journal, 2007, 6, 298–305.

46 A. Klute and C. Dirksen, in Methods of Soil Analysis, John Wiley & Sons, Ltd, 1986, pp. 687–734.

47 E. E. Nuccio, E. Starr, U. Karaoz, E. L. Brodie, J. Zhou, S. G. Tringe, R. R. Malmstrom, T. Woyke, J. F. Banfield, M. K. Firestone and J. Pett-Ridge, ISME J, 2020, 14, 999–1014.

48 S. Ge, X. Dong, Y. Liu, K. M. Wright, S. N. Humphris, L. X. Dupuy and M. P. MacDonald, Journal of Experimental Botany, 2023, 74, 787–799.

